# CytoVerse: Single-Cell AI Foundation Models in the Browser

**DOI:** 10.64898/2026.01.29.702554

**Authors:** Robert Currie, Jesus Gonzalez-Ferrer, Mohammed A. Mostajo-Radji, David Haussler

## Abstract

Mapping single-cell datasets to large atlases is often hindered by server constraints and privacy concerns. We present CytoVerse, a framework that runs scRNA-seq Foundation Models (scFM) entirely in the browser. Three key contributions enable this: (1) deploying models via ONNX without server side compute; (2) using compressed indexing (IVFPQ) to search a more then 20 million cell reference from the client; and (3) a lightweight protocol for sharing embeddings across consortia without exposing raw data. CytoVerse thereby provides a scalable, privacy preserving framework for distributed single-cell analysis.

## 1 Background

Mapping single-cell transcriptomic datasets to curated reference atlases has become central to cell type annotation and interpretation in single-cell RNA sequencing (scRNA-seq) [1, 2]. While existing web-based platforms such as Azimuth [3], SIMS-Web Streamlit [4, 5], MapMyCells [6], and ArchMap [7] have expanded access to reference-based annotation, they remain tethered to a server-side paradigm. This architecture creates three bottlenecks: (1) Ecosystem lock-in tied to specific languages or pre-integrated references; (2) File size limits and asynchronous processing that interrupt iterative exploration; and (3) Data privacy concerns when uploading unpublished or sensitive data.

Azimuth, a widely used platform, provides a graphical interface for reference mapping within the Seurat framework, but is designed around pre-integrated references stored in a specialized format. This tight integration with the Seurat ecosystem can make it less straightforward to incorporate custom references or workflows developed outside of R.

SIMS-Web Streamlit and MapMyCells enable browser-based cell type classification using deep learning models, offering convenient access without local setup. However, these approaches are primarily based on supervised models and depend on server-side computation, which can affect interactivity and impose practical limits on dataset size, typically on the order of a few gigabytes.

ArchMap extends flexibility by supporting multiple mapping backbones, including scVI, scANVI, and scPoli, and by providing a no-code interface for mapping queries across heterogeneous references. While this broadens usability, analyses are still performed through cloud-based workflows with predefined limits on query size, and results are returned after remote processing. In practice, users upload capped datasets, await asynchronous processing, and download static results. This workflow interrupts the iterative process of exploration and discovery. Meanwhile, improvements in tissue culture and sequencing throughput have generated reference atlases containing tens of millions of cells [8], emphasizing the need for more scalable and responsive analytical solutions.

Current server-based architectures also prevent users from fine tuning or adapting reference models, a limitation that is increasingly problematic in perturbation studies. Genetic [9] and chemical perturbations [8] can produce novel cellular states that are absent from training data, forcing out-of-distribution states into predefined labels, reducing biological interpretability.

Limitations on existing visualization platforms suggest there is a need for web applications that support scalable, interactive, and collaborative analysis before stable cell type labels are established.

WebAssembly (WASM) has emerged as a key technology for achieving high performance computation directly within the browser. The application Kana [10] exemplifies this approach by performing end-to-end scRNA-seq workflows (normalization, dimensionality reduction, clustering, marker detection, and annotation) entirely on the user’s device without requiring backend infrastructure. Kana demonstrates that, through WASM and optimized C++ backends, datasets with hundreds of thousands of cells can be processed within minutes while maintaining interactivity, privacy, and zero installation overhead. As WASM integrates with GPU accelerated browser APIs such as WebGPU [11], this model provides a foundation for next generation platforms capable of high throughput and collaborative single-cell analysis at the scale demanded by perturbation driven discovery.

Recently developed unsupervised scRNA-seq Foundation Models (scFM) [12, 13] function similarly to large language models, utilizing self supervised learning on vast, heterogeneous public repositories to capture universal representations of cellular biology. By addressing persistent challenges in annotation, nomenclature, and batch correction, they enable robust, ontology aware comparisons across multi-study atlases. Since these models require no retraining or metadata harmonization, deploying them via web applications is the logical next step to facilitate cross consortia collaboration within a shared latent space.

Building on these advances we present CytoVerse, a platform that shifts scFM from server pipelines to browser native workflows via: (1) Deployment of high parameter foundation models (e.g., SCimilarity [14]) directly in the browser via ONNX, bypassing the need for standard installed scRNA-seq software environments; (2) Application of Inverted File with Product Quantization (IVFPQ) [15] - a compression technique that reduces index size while preserving search accuracy - to enable high-throughput reference mapping against more then 20 million cells without incurring server-side computational costs; and (3) Introduction of a lightweight latent space sharing mechanism that enables consortium scale collaboration without the privacy or bandwidth overhead of raw data transfer.

## 2 Results

### 2.1 Browser-based Workflow and Data Flow

In CytoVerse, a researcher can embed, label, visualize, and interpret a local scRNA-seq dataset entirely within the browser (Figure 1). By replacing standard server-side pythonic pipelines with ONNX execution and an IVFPQ based indexing strategy, the client can map over 5,000 cells per minute on recent consumer laptops (e.g., Apple M-series). Once a user selects a local .h5ad file, CytoVerse performs the following steps using only static assets retrieved from the server:

**Fig. 1.**
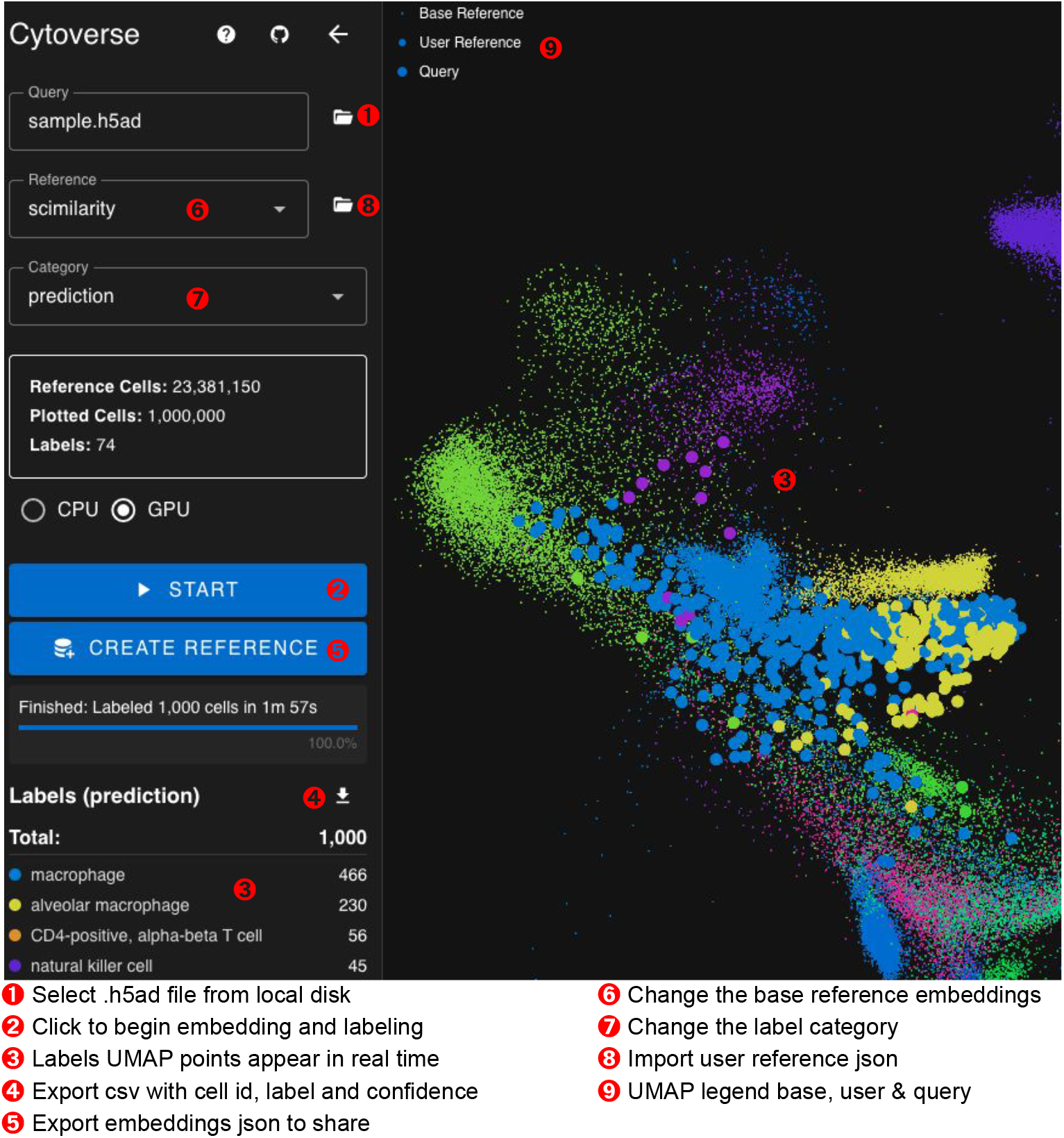
CytoVerse UI Guide

- Load and preprocess - Streams raw counts from a local .h5ad, normalizes, and aligns genes to the model’s expected input.
- Embed - Runs SCimilarity, or any ONNX-compatible model (e.g., SIMS [4]) to generate per-cell embeddings.
- Label - Assigns cell type labels using approximate nearest-neighbor search against a reference.
- Visualize - Projects embeddings into 2D via parametric UMAP [16] for interactive visualization and real-time plotting, including label-based coloring.
- Interpret - Computes a selected cell’s gene importance via embedding perturbation.
- Export - Export labels as CSV for downstream analysis and embeddings as JSON for latent space sharing.

### 2.2 Server Static Computation and Data Flow

The server’s role is limited to serving static files - it performs no computation at query time (Figure 2). During reference construction (a one time offline step), embeddings are partitioned into 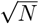 clusters via k-means, each stored as a binary file with PQ-compressed vectors and cell labels. A centroid index enables efficient lookup. Once these static artifacts are generated, the cloud or server has no further computational burden: it simply serves the static data artifact files (partition binaries and centroid index) and the small ONNX model files over plain HTTP/HTTPS. This lightweight serving scales cost effectively via CDNs or object storage (e.g., S3).

**Fig. 2.**
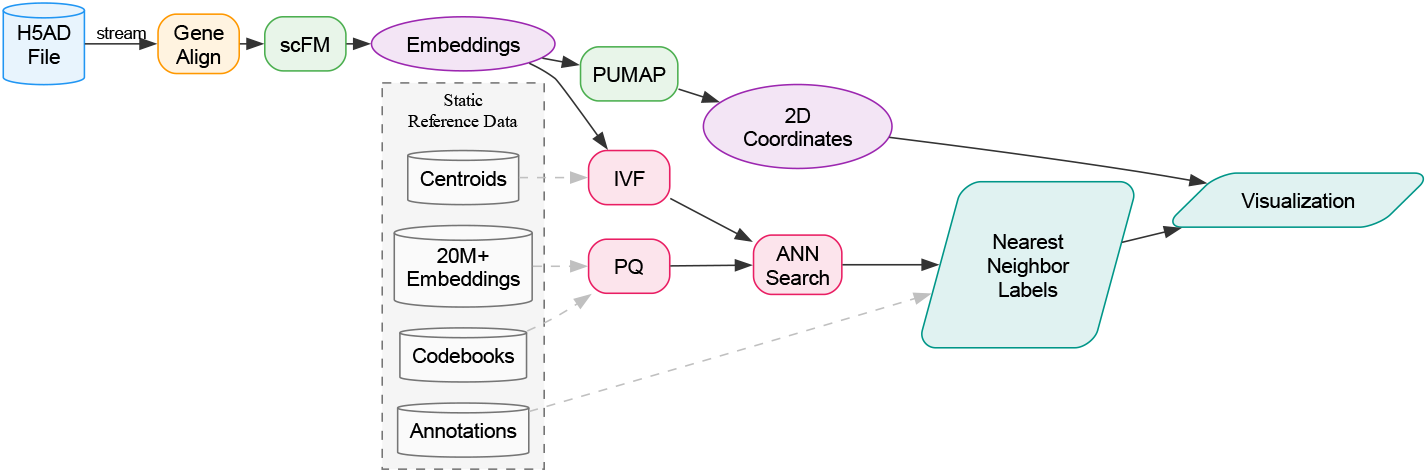
Compute & data flow for server statics assets generation and browser client computation

### 2.3 Query vs. Base and User Reference

To ensure consistent nomenclature across distributed analyses, CytoVerse distinguishes between two reference tiers. The **base reference** serves as the immutable foundation, defining the embedding space, nearest-neighbor index, and global label ontology. Complementing this, a **user reference** is a lightweight, portable subset JSON file constructed by projecting a focused query dataset (e.g., a .h5ad of brain cells) into the base reference. Crucially, a user reference does not define new cell categories; rather, it acts as a curated window into the base reference, ensuring that all annotations remain grounded in the standardized vocabulary. Researchers can share these user references as compact, embedding only JSON files, enabling collaborative analysis in latent space while preserving the privacy of the underlying raw data.

### 2.4 Model Explainability

CytoVerse implements a perturbation based feature importance method [17]. For each cell, we compute a baseline embedding, then iteratively zero out each expressed gene (expression *>* 0.1) and measure the L2 distance between baseline and perturbed embeddings. Genes whose removal causes larger distortions receive higher importance scores.

This method successfully recovers cell type enriched genes consistent with known biology. For example, analyzing Inhibitory and Excitatory neurons reveals distinct and shared importance profiles (Figure 3, left and right panels). Both cells share KCNC2, a gene that encodes for the Kv3.2 channel, important for neuronal excitability [18]. Inhibitory neurons show high importance scores for genes involved in specific circuit formation and modulation, including the axon guidance receptor ROBO2 (importance = 0.0108) and the glutamatergic receptor subunit, GRIK1 receptor (importance = 0.0103), that mediates excitatory neurotransmission in interneurons [19, 20]. In contrast, Excitatory neurons prioritize the LDL receptor related protein LRP1B (importance= 0.0064) and the RNA binding protein RALYL (importance = 0.0057), both enriched in excitatory populations [21, 22]. While some high expression genes like MALAT1 appear in both lists, their differential ranking and the emergence of cell type enriched markers like GRIK1 and LRP1B demonstrate that the model captures the distinct molecular identities defining these major neuronal classes.

**Fig. 3.**
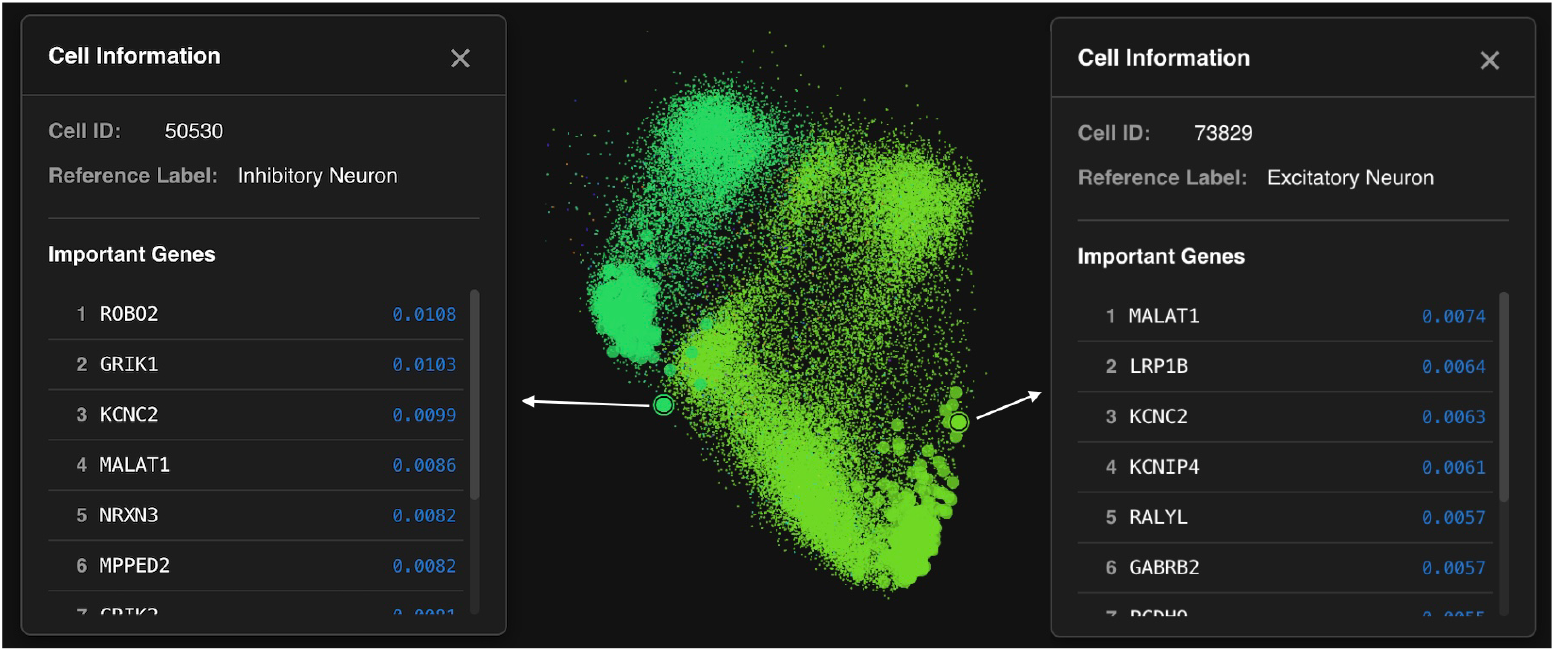
Important gene’s driving differences in embeddings between inhibitory and excitatory neurons against the SSPsyGene reference

### 2.5 Use Case: Data sharing across the SSPsyGene Consortium

Any .h5ad expression file can be used to generate a base reference hosted on the server and shared with all users. Alternatively, individual researchers can export embeddings and labels from their own datasets to share as ad hoc user references, facilitating iterative comparison and collaboration within a consortium. To illustrate these use cases we used an atlas of 600,000 fetal brain cells from the Nano *et al*., 2025 dataset [23]. This dataset (Figure 4) was selected as the primary reference for comparing data across developmental human brain models within the SSPsyGene [24] consortium, serving as a domain specific foundation for cross model alignment. Members can label results against SSPsyGene and share as user references for ad hoc collaboration. To evaluate performance, we collected datasets produced by a member of the consortium[25] and classified 10,000 NGN2 neurons against the SSPsyGene reference, achieving *>*80% accuracy in under a minute with most misclassifications assigned to inhibitory neurons and showing markers such as DLX6-AS1[26] (Figure 4).

**Fig. 4.**
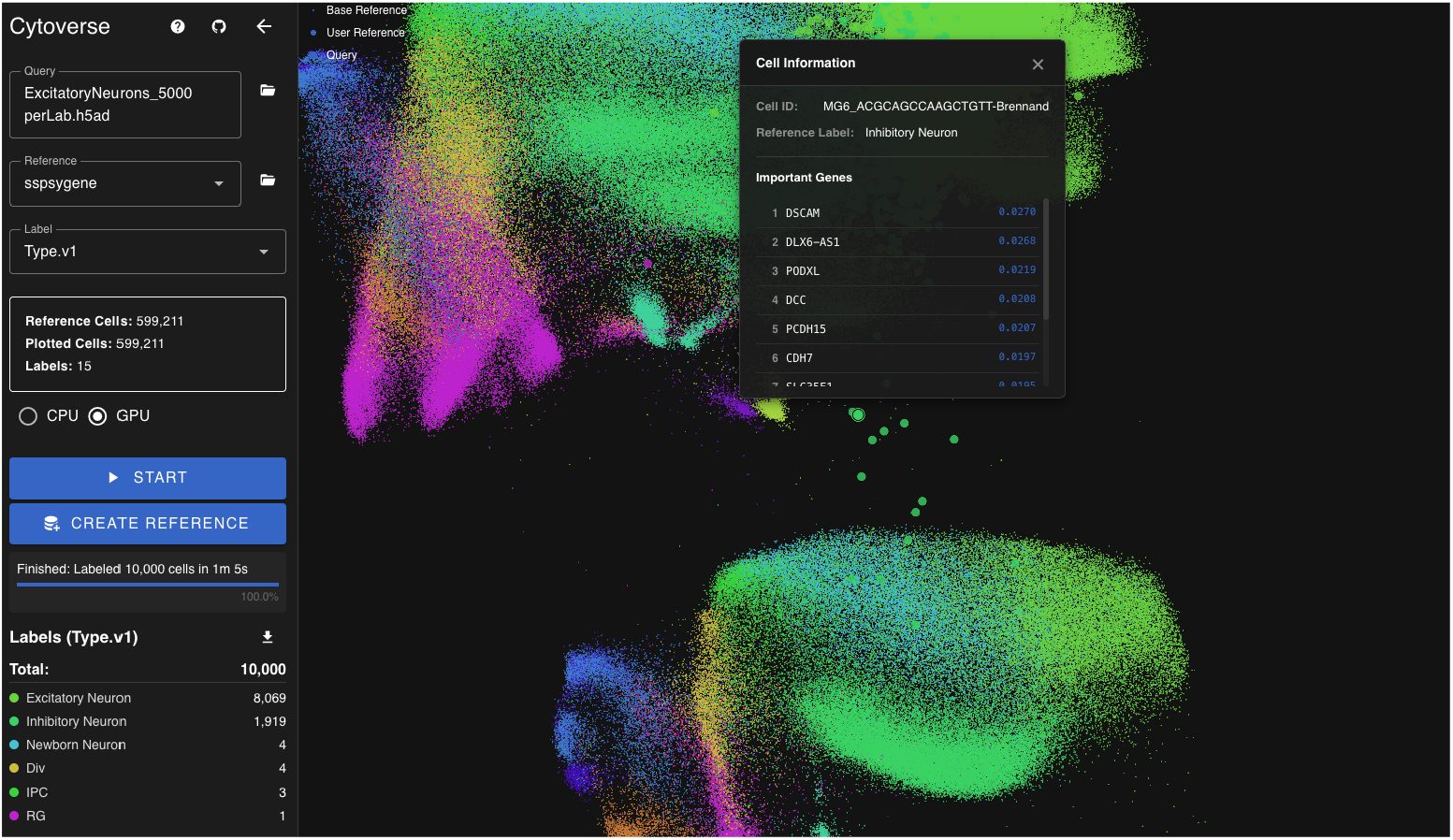
NGN2 neurons labeled against the SSPsyGene reference

### 2.6 Concordance

CytoVerse prioritizes rendering foundation model analyses accessible via interactive, browser based workflows with zero server compute. This design necessitates specific trade offs between fidelity and efficiency, most notably through the use of IVFPQ. While the underlying IVF structure and lossy PQ compression introduce an expected recall gap compared to exact server-side search, they are essential for operating within browser constraints. This compromise enables high throughput client-side performance facilitating immediate feedback and lightweight collaboration at the point of assay generation. Consequently, we position CytoVerse not as a replacement for final, resource intensive server pipelines, but as an accelerator for iterative discovery, course correction lightweight latent space collaboration close to assay generation.

#### 2.6.1 Concordance - Embeddings

Client side computation may require model ablation to ensure compatibility with browser-based environments. Such reductions can potentially affect model fidelity and reproducibility. To verify that the browser deployed model maintains equivalence to the original PyTorch implementation, we embedded 100 randomly selected expression profiles from the holdout dataset GSE136831.h5ad. The maximum embedding difference between the original SCimilarity model in PyTorch and its ONNX export evaluated with the ONNX Runtime was 3.31 × 10^−6^, indicating near perfect concordance. In parallel we validated similar concordance to five decimal places with the SIMS Web ONNX implementation.

#### 2.6.2 Concordance - Labels

Exact K-Nearest Neighbor (KNN) search requires a 20 GB SCimilarity index. This overwhelms standard client devices. IVFPQ resolves this by conceptually extending the logic of representation learning itself: just as embeddings compress gene counts to capture the intrinsic biological manifold, IVFPQ further compresses those embeddings by discarding redundant floating point precision. We significantly reduce bandwidth and memory load by transmitting only the structural signal required to identify cell states, rather than the “full resolution” noise. By varying the number of subspaces and probe count we can tradeoff the mismatch rate with bandwidth and client memory (Table 1). While this introduces a controlled decrease in recall (Figure 5) and distortion of the encoded latent space (Figure 6), the essential biological relationships remain intact (Table 2).

**Table 1.** Comparative analysis of label changes compared to full KNN as IVFPQ parameters are increased from low to high (used in the deployed application)

| Metric | Lowest Parameters<br>($n_{\text{sub}} = 8, n_{\text{probe}} = 1$) | Highest Parameters<br>($n_{\text{sub}} = 32, n_{\text{probe}} = 32$) |
| --- | --- | --- |
| Mismatch Rate | 47.0% | 13.0% |
| Total Mismatches | 47 | 13 |
| Improvement | – | 34.0 pp |

**Table 2.**
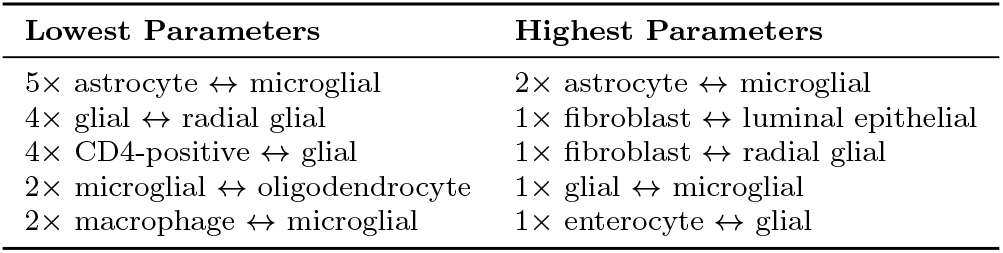
Examples of label changes as IVFPQ compression increases demonstrating essential biological structure is largely preserved.

| Lowest Parameters | Highest Parameters |
| --- | --- |
| 5× astrocyte ↔ microglial | 2× astrocyte ↔ microglial |
| 4× glial ↔ radial glial | 1× fibroblast ↔ luminal epithelial |
| 4× CD4-positive ↔ glial | 1× fibroblast ↔ radial glial |
| 2× microglial ↔ oligodendrocyte | 1× glial ↔ microglial |
| 2× macrophage ↔ microglial | 1× enterocyte ↔ glial |

**Fig. 5.**
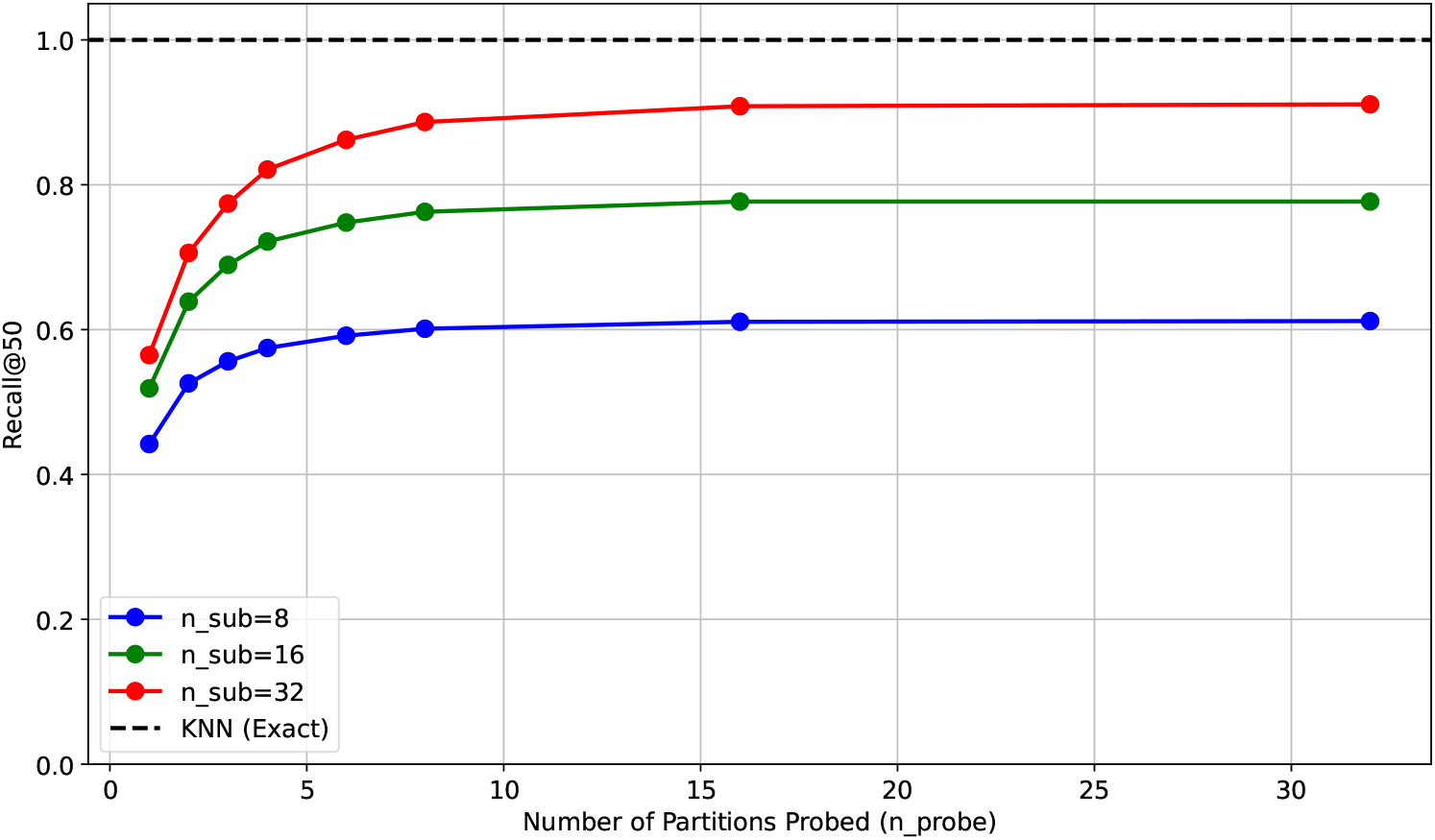
Exact kNN baseline shown as a dashed line vs. IVFPQ recall as the number of IVFPQ partitions probed (bottom axis) and the number of PQ subspaces (red, green and blue curves) increase.

**Fig. 6.**
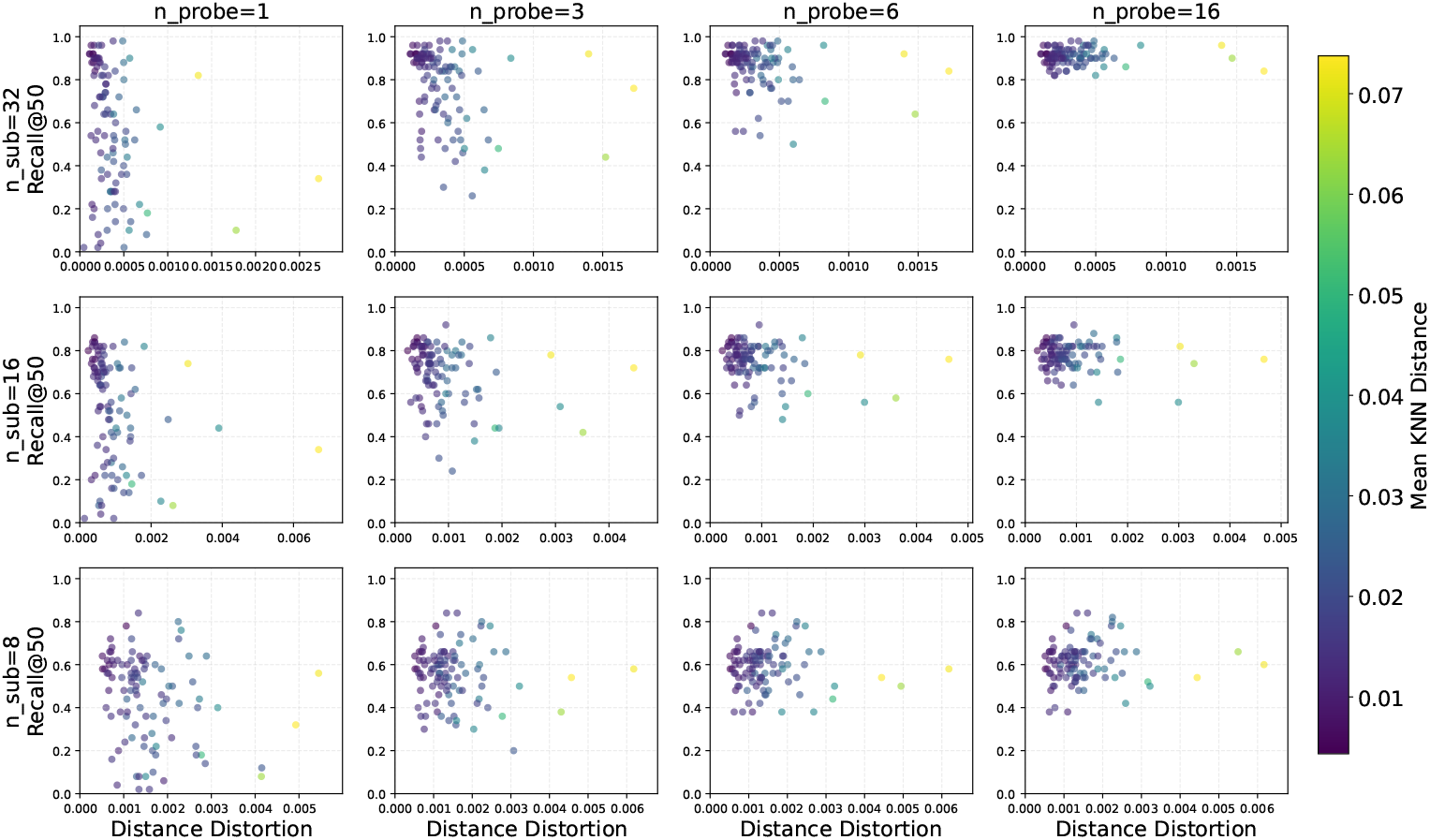
Latent space distance distortion vs. exact kNN as probed IVF partitions increase left to right and PQ subspaces increase bottom to top.

### 2.7 Generalization to other models

Any embedding model exportable to ONNX can be integrated. PyTorch models use torch.onnx.export; TensorFlow models use tf2onnx.convert. We previously adapted SIMS and parametric UMAP for browser execution, informing the current design. New models can be incorporated as they are validated.

## 3 Discussion

CytoVerse introduces a technical framework for single-cell analysis that allows client-side execution not possible in traditional cloud-based architectures. Our work demonstrates that the inherent barriers to browser-based AI can be overcome through ONNX-based deployment and IVFPQ-enabled reference mapping. By eliminating the reliance on backend infrastructure and expensive server-side compute, CytoVerse makes high-throughput foundation model analysis economically accessible at the point of assay generation.

This technical shift directly addresses the sociological and privacy-related bottlenecks inherent in modern functional genomics. Scientific discovery is fundamentally collaborative, yet the practicalities of sharing raw data often result in significant delays, with data exchange frequently occurring only upon formal publication. Within consortia, these delays limit early opportunities for cross-laboratory comparison, redundancy detection, and joint interpretation of emerging datasets. CytoVerse facilitates collaboration directly in latent space by enabling the export and exchange of embeddings and labels as lightweight JSON files. This framework allows researchers to share curated “windows” into a base reference, ensuring that all annotations remain grounded in a standardized vocabulary while preserving the privacy of the underlying raw data.

As foundation models and browser technologies continue to advance, CytoVerse provides an extensible architecture for integrating new embedding models, multi-modal datasets, and advanced visualization tools. In the long term, it lays the groundwork for a future where single-cell atlases are no longer static, centralized repositories, but are instead continuously refined, queried, and interpreted in real-time by distributed scientific communities.

## 4 Methods

The following sections summarize the core components and enabling technologies that allow all computations to be executed directly within the web browser.

### 4.1 Expression File Access and Preprocessing

AnnData [27] .`h5ad` files are read client-side using `h5wasm` (https://github.com/usnistgov/h5wasm), a zero dependency WebAssembly based library for streaming local HDF5 files. The library is implemented directly on the native HDF5 C API, ensuring full compatibility with standard HDF5 datasets and avoiding translation layers that can compromise performance or fidelity.

### 4.2 Computation

ONNX [28] defines an extensible, hardware agnostic computation graph that includes standardized operators, data types, and parameters. Within the browser, ONNX runtimes support both CPU and GPU execution, allowing .onnx models to run with near native performance. In CytoVerse, ONNX models perform key analytical steps, including count and Lp-norm preprocessing of gene expression data, expression embedding, centroid lookup, product quantization (PQ)–based asymmetric distance ranking, and parametric UMAP projection. Equivalent server side ONNX runtimes enable the same .onnx models to be executed at scale for reference population, ensuring precise consistency and reproducibility without reliance on containerized workflows.

### 4.3 Labeling

The IVFPQ database partitions the reference dataset into approximately 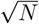 clusters, where *N* represents the total number of cells or embeddings. Each partition is stored on the server and retrieved on demand through HTTP requests initiated by centroid based search. For the SCimilarity reference set comprising 23.3 million cells, the median partition size is 130 KB. Because query datasets are typically homogeneous, only a small subset of partitions needs to be fetched; these are retrieved in parallel and cached locally during each labeling operation.

### 4.4 User Interface

The CytoVerse user interface is implemented in the Vue (https://vuejs.org/) TypeScript framework and utilizes browser Web Workers to parallelize computation, maintaining responsiveness during analysis. A parametric UMAP model [16] is trained using `umap-pytorch` on a stratified subset of embeddings and exported to ONNX format. During runtime, this model generates two dimensional coordinates for labeled cells in real time as the query file is streamed and embedded. The resulting points are visualized using `regl-scatterplot` (https://github.com/flekschas/regl-scatterplot), which employs a high performance WebGL shader engine to render more than one million points interactively with smooth navigation and dynamic label coloring.

### 4.5 Datasets

SCimilarity pretrained model weights and embeddings were downloaded from: https://zenodo.org/records/10685499

SCimilarity human tissue atlas scRNA-seq data with metadata and can be downloaded from: https://zenodo.org/records/10895214

The SSPsyGene consortium reference dataset was based on the Integrated development Metaatlas [23]

The Human Cortex M1 region dataset from the Allen Institute [29] was obtained via the UCSC Cell Browser.

The dataset used to evaluate the SSPsyGene consortium reference was compiled from previously published studies generated by laboratories within the consortium [25].

## 4.6 Acknowledgements

## 4.7 Authors’ contributions

R.C. and J.G.-F. conceived the project and performed the experiments. M.A.M.-R. and D.H. supervised the work and secured the funding. All others wrote the manuscript.

The authors would like to acknowledge the assistance of the Claude, Gemini and Grok large language models for generating portions of the source code, various UI elements and analysis figures. The authors retain full responsibility for all content generated by the AI.

## 4.9 Availability of data and materials

CytoVerse is accessible at https://cells-test.gi.ucsc.edu/cytoverse/, and all source code and tools are available at https://github.com/braingeneers/cytoverse.

## Declarations

### 4.10 Ethics approval and consent to participate

Not applicable.

### 4.11 Consent for publication

Not applicable.

